# Flexible recruitments of fundamental muscle synergies in the trunk and lower limbs for highly variable movements and postures

**DOI:** 10.1101/2021.08.03.455001

**Authors:** Hiroki Saito, Hikaru Yokoyama, Atsushi Sasaki, Tatsuya Kato, Kimitaka Nakazawa

**Author notes:** **Corresponding authors**: Kimitaka Nakazawa, PhD, The University of Tokyo, Department of Life Sciences, Graduate School of Arts and Sciences, 3-8-1 Komaba, Meguro-ku, Tokyo, 153-8902, Japan.

## Abstract

The extent to which muscle synergies represent the neural control of human behavior remains unknown. Here, we tested whether certain sets of muscle synergies that are fundamentally necessary across behaviors exist. We measured the electromyographic activities of 26 muscles including bilateral trunk and lower limb muscles during 24 locomotion, dynamic and static stability tasks, and extracted the muscle synergies using non-negative matrix factorization. Our results showed that 13 muscle synergies that may have unique functional roles accounted for almost all 24 tasks by combinations of single and/or merging of synergies. Therefore, our results may support the notion of the low dimensionality in motor outputs, in which the central nervous system flexibly recruits fundamental muscle synergies to execute diverse human behaviors. Further studies using manipulations of the central nervous system and/or neural recording are required the neural representation with such fundamental components of muscle synergies.

## Introduction

To execute human movements, the central nervous system (CNS) must control many degrees of freedom from thousands of motor units within hundreds of skeletal muscles ^1^. To simplify the production of movements, the CNS may rely on a limited number of neural mechanisms ^2^. Indeed, the CNS exploits a reduced set of pre-shaped neural pathways, called muscle synergies, to achieve a large variety of motor commands ^3,4^. Muscle synergy theory assumes that the CNS combines a few sets of activation to build muscle activation commands ^5^. Evidence of a limited set of muscle synergies has been found in various human motor behaviors such as locomotion ^6–9^, reaching tasks ^10^, and sports activities ^11–13^.

It has been proposed that muscle synergies are shared across various motor tasks ^5,14,15^. Shared synergies facilitate the robustness of the neuromuscular system, which is thought to be beneficial for stable postural control ^15,16^, development ^17^, and expert motor skills ^18^. In contrast, studies have also discovered the existence of task specific synergies to meet each biomechanical demand of motor tasks ^19,20^. An experimental study in frogs investigated muscle synergies during natural behaviors such as walking, jumping, and swimming, indicating that each motor behavior is the consequence of a combination of both synergies shared between behaviors and synergies specific to each or a few behaviors ^21^. However, a substantial number of in-born and learned human movements and postures that presents different behavioral contexts also exist ^22^. As such, it is possible that the sum of all shared and task-specific synergies employed during a variety of human movements and postures may exceed the number of relevant muscles ^23^, violating the existence of a low dimensionality of human movement controls based on muscle synergy theory ^23,24^.

A previous study found that upper-limb hand exploration tasks for five sectors (frontal, right, left, horizontal, and up) were modulated by seven muscle synergies (i.e., seven cluster centroids across participants) with different functional roles ^14^. Furthermore, another study found that all three muscle synergies of cycling can be well reconstructed by merging muscle synergies extracted from walking ^25^. Thus, the interpretation of existing literature suggests that the CNS may select the appropriate subsets of muscle synergies, either independently or merged, from a large set that are established to execute the substantial number of behavioral contexts and demands ^14,25,26^. However, previous studies have not recorded a large set of electromyographic (EMG) activities during a variety of human movements and postural tasks with different biomechanical contexts to investigate the neural basis of muscle synergies.

We hypothesized the existence of fundamentally necessary muscle synergies that account for a diverse range of human movements and postures. To investigate this possibility, we first extracted muscle synergies from an EMG recording dataset made from 24 motor tasks of the trunk and lower limb muscles to define the fundamental muscle synergies utilized across a highly variable context of movements and postures. We then examined how these fundamental muscle synergies were used in each motor task by comparing them to those extracted from the EMG datasets in each task.

## Methods

### Experimental protocol

Ten healthy volunteers (aged 21−35 years, all men) participated in the study. Each participant provided written informed consent for participation in the study. The study was conducted in accordance with the Declaration of Helsinki and was approved by the local ethics committee of the University of Tokyo.

We focused on fundamental movement and postural tasks that serve as building blocks for the efficient and effective execution of a variety of daily living activities and highly skilled actions such as sports ^22,27,28^. Specifically, we used tasks that required movements through space (locomotion) and controls against gravity (stability) in any plane ^22^. Thus, all participants were asked to perform the 24 tasks described in Table 1. Supplementary Table S1 online presents the details of each movement and postural task. The order of tasks was randomly assigned.

**Table 1.**
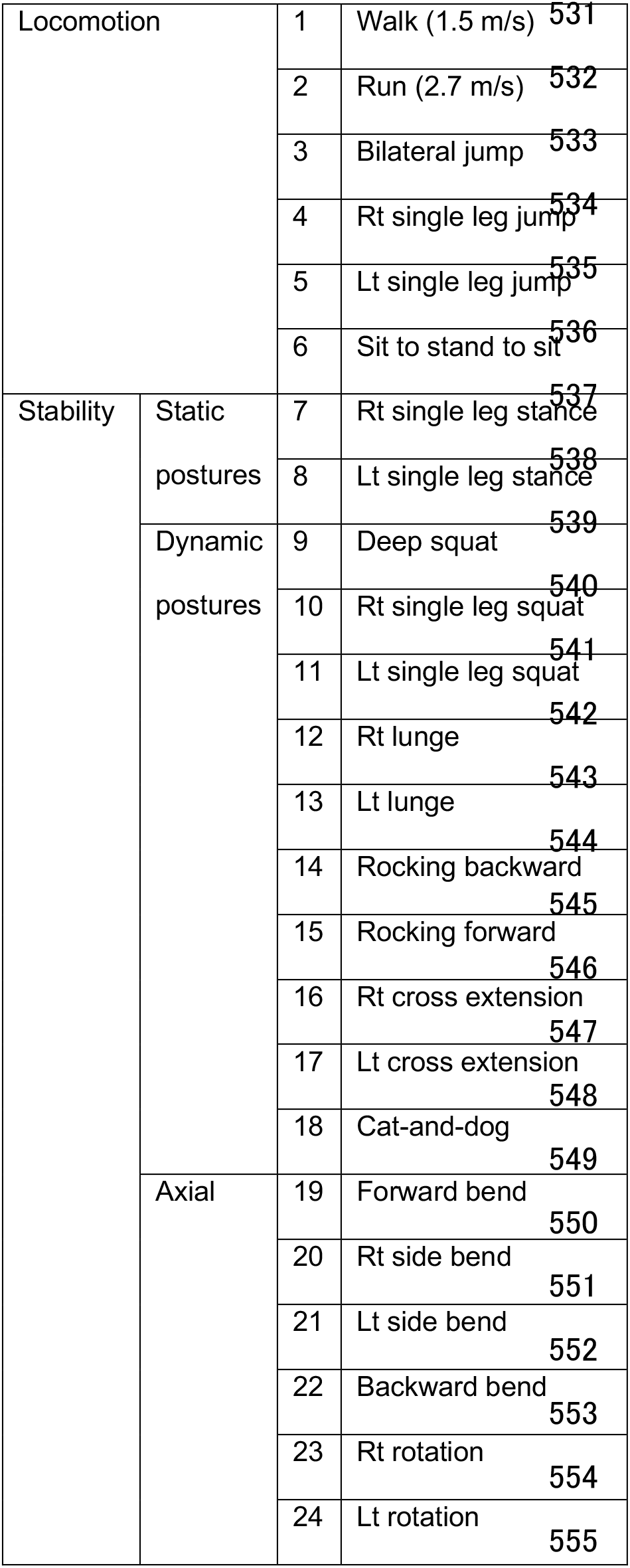
Movement and postural tasks. Shown are the order of 24 locomotion and stability task. Stability tasks are divided into three subcategories: static postures, dynamic postures and axial. Rt: right; Lt: left.

|  |  |  |  |  |
| --- | --- | --- | --- | --- |
| Locomotion |  | 1 | Walk (1.5 m/s) | 531 |
|  |  | 2 | Run (2.7 m/s) | 532 |
|  |  | 3 | Bilateral jump | 533 |
|  |  | 4 | Rt single leg jump | 534 |
|  |  | 5 | Lt single leg jump | 535 |
|  |  | 6 | Sit to stand to sit | 536 |
| Stability | Static postures | 7 | Rt single leg stance | 537 |
|  |  | 8 | Lt single leg stance | 538 |
|  | Dynamic postures | 9 | Deep squat | 539 |
|  |  | 10 | Rt single leg squat | 540 |
|  |  | 11 | Lt single leg squat | 541 |
|  |  | 12 | Rt lunge | 542 |
|  |  | 13 | Lt lunge | 543 |
|  |  | 14 | Rocking backward | 544 |
|  |  | 15 | Rocking forward | 545 |
|  |  | 16 | Rt cross extension | 546 |
|  |  | 17 | Lt cross extension | 547 |
|  |  | 18 | Cat-and-dog | 548 |
|  | Axial | 19 | Forward bend | 549 |
|  |  | 20 | Rt side bend | 550 |
|  |  | 21 | Lt side bend | 551 |
|  |  | 22 | Backward bend | 552 |
|  |  | 23 | Rt rotation | 553 |
|  |  | 24 | Lt rotation | 554 |
|  |  |  |  | 555 |

### Data collection

EMG activity was recorded from the following 26 muscles distributed across the trunk and lower limbs (13 bilateral muscles): tibialis anterior (TA), gastrocnemius medialis (MG), vastus medialis (VM), rectus femoris (RF), biceps femoris (long head, BF), gluteus maximus (GM), gluteus medius (Gmed), rectus abdominis (RA), oblique externus (OE), erector spinae at L2 (ESL2), erector spinae at Th9 (ESTh9), erector spinae at Th1 (ESTh1), and latissimus dorsi (LD). EMG activity was recorded using a wireless EMG system (Trigno Wireless System; DELSYS, Boston, MA, USA). The EMG signals were bandpass filtered (20–450 Hz), amplified (with a 300-gain preamplifier), and sampled at 1000 Hz. Three-dimensional ground reaction force data were recorded at 1000 Hz from the force plates under each belt of the treadmill.

### EMG processing

The low-pass cut-off frequency influences the smoothing of EMG patterns and thus impacts the number of extracted modules ^29^. To adequately compare EMG envelopes (i.e., EMG patterns with the same smoothing) of movements performed for various tasks that had different features of dynamic activities, the low-pass cut-off frequency must be adjusted for each task. Thus, an iterative adaptive algorithm was used to extract the optimal EMG envelopes ^30^. This algorithm utilized information theory to find a sample-by-sample optimal root-mean-square window for envelope estimation ^30^. This algorithm allowed the filter to adequately follow fast changes in EMG activity while maintaining optimal extraction when the EMG amplitude is changing slowly ^30^. A previous study used this algorithm and successfully reconstructed muscle synergies during walking in individuals with and without transfemoral amputation ^31^. The smoothed EMG envelopes were time-interpolated to generate 200 timepoints for each trial, except for the right and left single-leg stance tasks.

We created the following two types of EMG matrices for each subject to examine the repertoire of fundamentally necessary muscle synergies and how these synergies are used in each task. Similar to previous studies ^14,32^, we pooled the EMG matrices of all 24 tasks to create an “all-task” EMG matrix for each subject (i.e., the matrix was composed of the 26 muscles × the summation of timepoints of the 24 single-task EMG matrices) to extract fundamental muscle synergies across all tasks. We also created a “single-task” EMG matrix composed of the 26 muscles × 1400 timepoints (seven strides or repetitions × 200 timepoints for each task, except the right and left single-leg stances) for each of the 24 tasks to extract muscle synergies.

### Muscle synergy analysis

In our analysis, we first identified the muscle synergies of each task for each subject using a factorization algorithm of single-task EMG matrices, and then synergies of all tasks were extracted from all-task EMG matrices using the same algorithm. We then proceeded to characterize representative muscle synergies of individual tasks and all tasks across all participants using a hierarchical clustering algorithm. Lastly, we analyzed the similarity between synergy cluster centroids of each individual task and single or merged synergies of the all-task matrix to investigate how muscle synergies utilized by all tasks contribute to the execution of each individual movement.

To explore muscle synergies, nonnegative matrix factorization (NMF) was used for each subject from the single-task EMG matrices and the all-task EMG matrix. NMF has previously been described as a linear decomposition technique ^33,34^ according to equation (1):

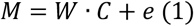

where *M* (*m* × *t* matrix, where *m* is the number of muscles and *t* is the number of samples, i.e., the spatiotemporal profiles of muscle activity) is a linear combination of muscle weighting components: *W* (*m* × *n* matrix, where n is the number of muscle synergies) and *C* (*n* × *t* matrix, representing temporal pattern components; and *e* is the residual error matrix. Each EMG vector in the matrix corresponding to each muscle activity was normalized to the maximum amplitude across all tasks so that all muscle scales ranged from 0 to 1. Prior to extracting muscle synergies, each muscle vector in the data matrix was standardized to have unit variance, thus ensuring that the activity in all muscles was equally weighted. However, after each synergy extraction, the unit variance scaling was removed from the data so that each muscle variable ranged from 0 to 1 for data inspection and interpretation ^35^.To determine the number of muscle synergies, NMF was applied to extract each possible *n* from 1 to 26 from each dataset. The variance accounted for (VAF) by the reconstructed EMG (*M*) was calculated at each iteration to extract the optimal number of muscle synergies. VAF was defined as a 100 × square of the uncentered Pearson’s correlation coefficient ^36,37^. To prevent the extracted synergies from assuming a suboptimal local minimum, each synergy extraction was repeated 100 times. Thus, the iteration with the highest VAF was maintained ^8^. We defined the optimal number *n* as the number fulfilling the following two criteria: First, *n* was selected as the smallest number of modules that accounted for >90% of the VAF ^36^. Second, *n* was the smallest number to which adding another module did not increase VAF by >5% ^38^.

### Clustering the modules across participants

We identified the representative synergy vectors across participants using hierarchical clustering analysis (Ward’s method, Euclidian distance) of muscle synergies for each task and all tasks ^8,39^. The optimal number of clusters was determined using the gap statistic ^40^. Subsequently, the muscle synergies in each cluster were averaged across participants.

### Contributions of the muscle synergy of all tasks to the execution of each task

To explore whether the muscle synergy defined by the all-task matrix contributes to executing each task of movements and postures, the similarity between muscle synergies of single-task and all-task matrices was quantified by the scalar product (SP) between these centroids of the synergy clusters (normalized to unit vectors). For every comparison, each of the synergy cluster centroids of all-task was matched to a synergy cluster centroid of each task by maximizing the total scalar product values. Synergy clusters that could not be matched with SP ≥0.75, were classified as unmatched ^41^.

### Contributions of merging muscle synergy of all tasks towards single-task execution

We also expected that all-task muscle synergies can be merged to execute each single task of movement and posture ^40^. Thus, the merged synergies as a linear combination of the contributing synergies were modeled by the following formula ^18,41^:

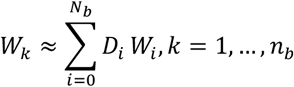

where ***W**_k_* is the *k*th muscle synergy vector from each individual task, ***W**_i_* is the *i*th muscle synergy vector derived from an all-task matrix, ***N**_b_* is the number of synergies that contribute to the merging, and ***D**_i_* is a non-negative coefficient that scales the *i*th synergy in the merging. ***D**_i_* was obtained from a non-negative least-squares fit, implemented using MATLAB (function lsqnonneg). ***W**_k_* and ***W**_i_* were normalized as unit vectors. Following criteria from previous studies ^18,41^, the synergy merging was identified when ***N**_b_* ≥ 2, ***D**_i_* ≥0.2 for all *i*, and the SP between 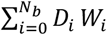 and ***W**_k_* was ≥0.75.

To assess whether the synergies from each task can be explained as merging of synergies from all tasks, we first identified the synergy cluster centroids of single-task and synergies of the all-task (described above) and reconstructed each synergy cluster centroid of each individual task by merging every possible combination of the synergy cluster centroids of all tasks.

## Results

### Muscle synergies extracted from all-task EMG matrices

Figure 1 presents 13 muscle synergies of an all-task matrix incorporating 24 trunk and lower limb movement tasks (W1 to W13), which were grouped by cluster analysis across ten participants; Table 2 summarizes the characteristics of the muscle synergies. Visual inspection revealed that muscle synergies W1 to W5 were largely composed of the right-side muscles, while muscle synergies W6 to W10 were mainly composed of the left-side muscles. Thus, we categorized W1 to W5 as muscle synergies with right-side dominant patterns and W6 to W10 as muscle synergies with left-side dominant patterns. The following pairs showed high similarity when the muscles in W6 to W10 were reordered so that muscles on the left side of W6 to W10 corresponded to the same muscles on the right side of W1 to W5: W1 and W6 (SP = 0.93), W2 and W7 (SP = 0.97), W3 and W8 (SP = 0.97), W4 and W9 (SP = 0.85), W5, and W10 (SP = 0.93). Others such as W11, W12, and W13 were categorized as bilateral patterns.

**Table 2.** Characteristics of muscle synergy clusters of all tasks. The following pairs showed high similarity when the muscles in W6 to W10 were reordered so that muscles on the left side of W6 to W10 corresponded to the same muscles on the right side of W1 to W5: W1 and W6 (SP = 0.93), W2 and W7 (SP = 0.97), W3 and W8 (SP = 0.97), W4 and W9 (SP = 0.85), W5, and W10 (SP = 0.93). We categorized W1 to W5 as muscle synergies with right-side dominant patterns and W6 to W10 as muscle synergies with left-side dominant patterns. W11, W12, and W13 were categorized as bilateral patterns. Muscles that account for > 0.5 of activation levels are classified as major muscles and between 0.1 to 0.5 were as minor muscles. isp: ipsilateral, con: contralateral, bil: bilateral.

| Unilateral patterns |  | Major muscles | Minor muscles |
| --- | --- | --- | --- |
| Right patterns | Left patterns |  |  |
| W1 | W6 | ispTA, ispRF, ispVM | (ispESL2, ispEST9, ispEST1, conTA, conESL2, conEST9, conLD) |
| W2 | W7 | ispVM, ispRF, ispGM, ispGmed | (ispMG, ispOE, conBF, conOE, conESL2) |
| W3 | W8 | ispMG, ispGmed | (ispRF, ispVM, ispBF, ispGM, ispEST1, ispLD, conTA, conBF, conOE, conESL2) |
| W4 | W9 | ispBF | (ispMG, ispGM, ispOE, ispESL2, conESL2, conEST9) |
| W5 | W10 | ispEST9, ispLD | (ispOE, ispESL2, ispEST1, conBF, conGM, conGmed, conOE, conESL2, conEST9, conEST1, conLD) |
| Bilateral patterns |  |  |  |
| M11 |  | bilESL2 | (bilEST9, bilEST1) |
| M12 |  | bilEST1 | (bilLD) |
| M13 |  | bilRAS, bilOE |  |

**Figure 1.**
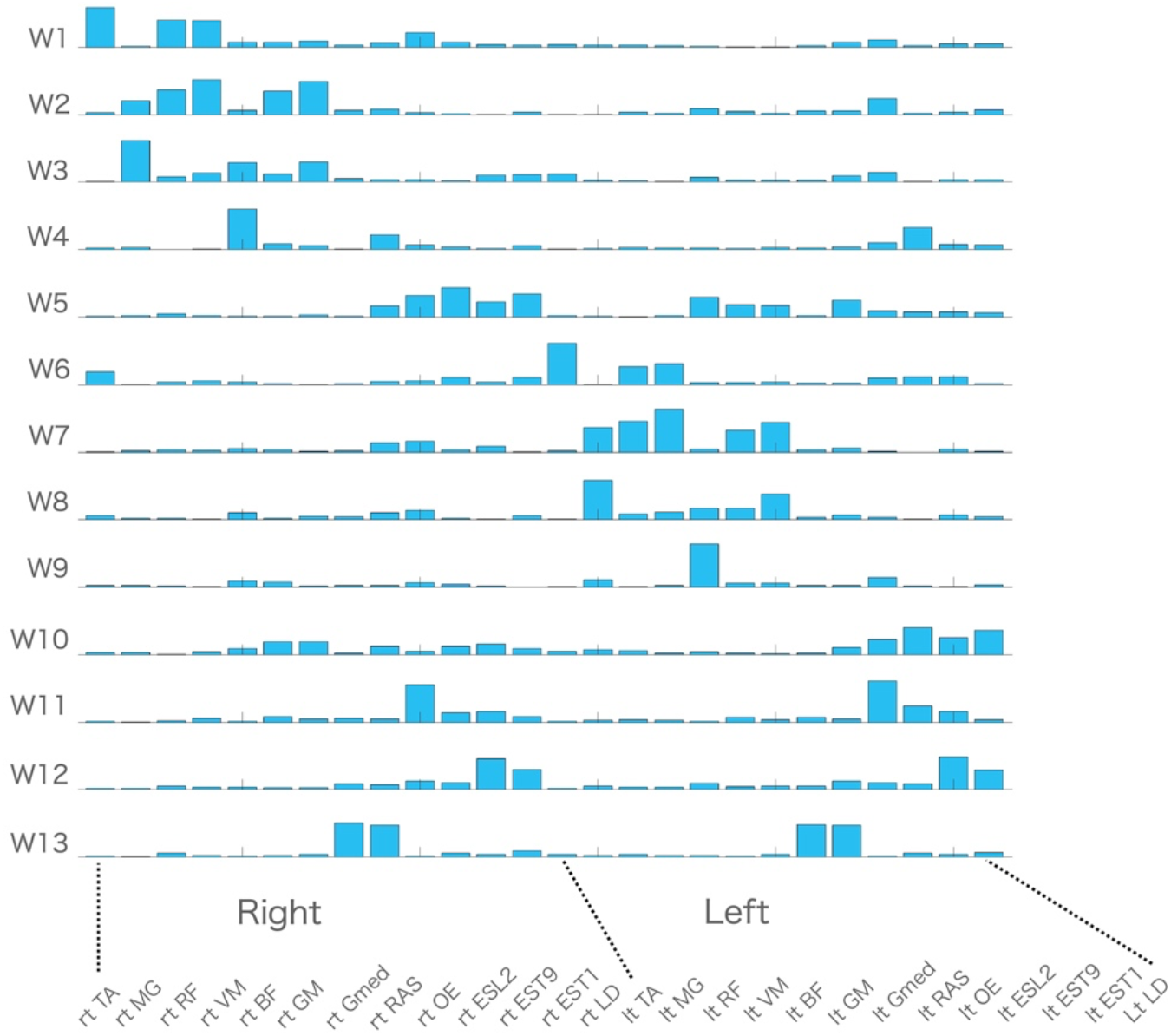
Muscle synergies of all tasks. **(a)** Centroids of the hierarchical clustering performed on the muscle synergies of all tasks across ten participants. **(b)** Dendrograms represent the results of cluster analysis (Ward’s method, Euclidian distance) where optimal number of clusters were determined based on the gap statistics.

### Relationship between muscle synergies extracted from all-task EMG matrices and those extracted from single-task matrices

Table 3 presents the number of muscle synergies in each task, which were well explained (SP > 0.75) by independent and merged muscle synergies from the all-task EMG matrices. Of note, all synergies of each task except the one for the left single-leg stance could be explained by either single or linear combination of multiple synergies from the all-task EMG matrices (SP > 0.75). The details of the contributions of muscle synergies of all tasks to each task execution are presented in Supplementary Table S2 online.

**Table 3.** The relationship between synergy clusters of each task and synergy clusters of all tasks. The number of synergy clusters for each task and the number of a single or merged synergy cluster centroids of all tasks that were well matched (scalar product > 0.75) or unmatched to a synergy cluster centroid of each task. Bil: bilateral; Rt: right; Lt left; JP: jump; SJP: single leg jump; STS: sit-to-stand-to-sit; SLS: single leg stance; DS: deep squat; SS: single leg squat; LG: lunge; RB: rocking backward; RF: rocking forward; CE: cross extension; CD: cat-and-dog; FB; forward bend; SB: side bend; BB: backward bend; RT: rotation.

| Movement and postural tasks | LtRT | RtRT | BB | LtSB | RtSB | FB | CD | LtCE | RtCE | Rf | Rb | LtLG | RtLG | LtSS | RtSS | Ds | LtSLS | RtSLS | STS | LtSJp | RtSJp | BilJp | Run | Walk |
| --- | --- | --- | --- | --- | --- | --- | --- | --- | --- | --- | --- | --- | --- | --- | --- | --- | --- | --- | --- | --- | --- | --- | --- | --- |
| Number of total synergy clusters | 2 | 2 | 2 | 2 | 2 | 2 | 3 | 2 | 2 | 2 | 2 | 2 | 2 | 2 | 2 | 2 | 2 | 2 | 2 | 3 | 3 | 4 | 4 | 4 |
| Number of synergy clusters that are well matched by a single synergy cluster of all tasks | 1 | 1 | 0 | 0 | 0 | 0 | 1 | 0 | 0 | 1 | 1 | 0 | 0 | 0 | 0 | 0 | 0 | 0 | 0 | 0 | 0 | 0 | 0 | 0 |
| Number of synergy clusters that are well matched by merging synergy clusters of all tasks | 1 | 1 | 2 | 2 | 2 | 2 | 2 | 2 | 2 | 1 | 1 | 2 | 2 | 2 | 2 | 2 | 1 | 2 | 2 | 3 | 3 | 4 | 4 | 4 |
| Number of synergy clusters that are unmatched by synergies of all tasks | 0 | 0 | 0 | 0 | 0 | 0 | 0 | 0 | 0 | 0 | 0 | 0 | 0 | 0 | 0 | 0 | 1 | 0 | 0 | 0 | 0 | 0 | 0 | 0 |

Figures 2 and 3 present examples of relationships between muscle synergies from the all-task EMG matrices and those from the single-task EMG matrices: locomotion tasks including walking, running, bilateral jump and sit-to-stand-to-sit (Fig. 2), and stability tasks including left lunge, cat-and-dog, forward bend, and left rotation (Fig. 3). The relationships between muscle synergies from the all-task EMG matrices and those from the other single-task EMG matrices are shown in Supplementary Figs. S1 and S2 online.

**Figure 2.**
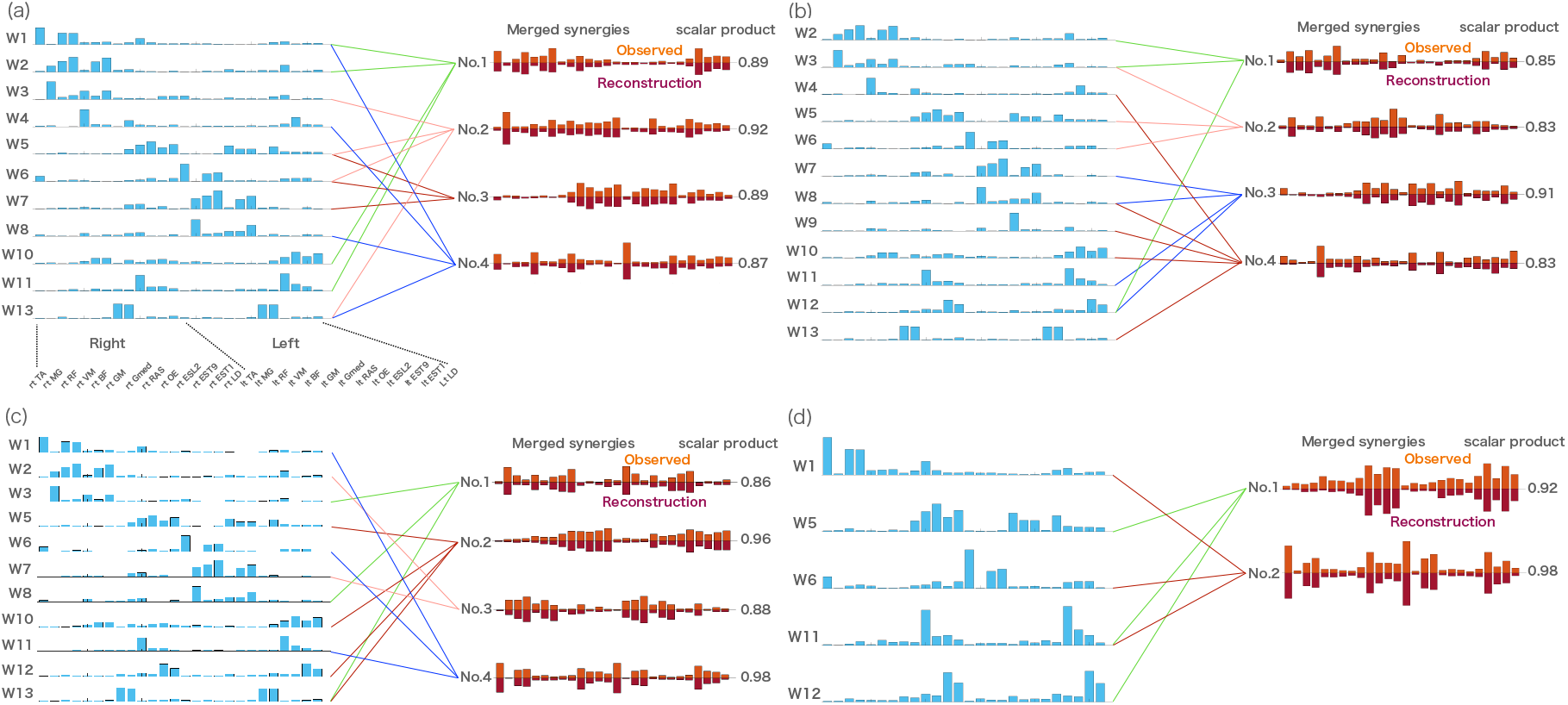
The relationship between muscle synergies of all tasks and muscle synergies of locomotion tasks including (a) walk, (b) run, (c) bilateral jump and (d) sit-to-stand-to-sit. The figures show the synergy cluster centroids of these tasks that could be explained by either a single or linearly combined multiple synergy cluster centroids of all tasks (synergies in blue) matched by maximizing scalar product > 0.75. Observed muscle synergies extracted from the single-task EMG (orange) and their reconstructions by merging their respective W1- combinations (dark orange) are further presented.

**Figure 3.**
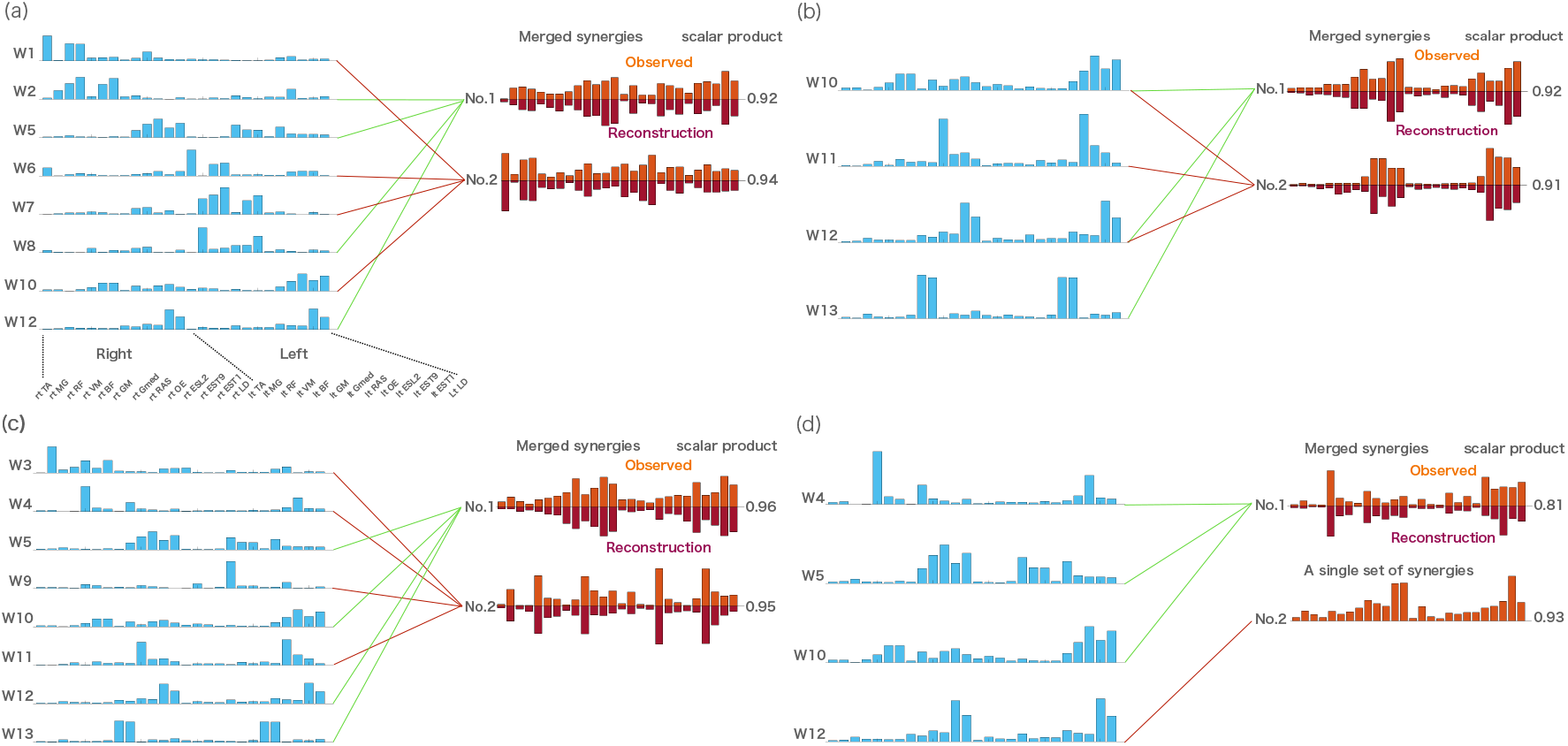
The relationship between muscle synergies of all tasks and muscle synergies of stability tasks including (a) left lunge, (b) cat-and-dog, (c) forward bend and (d) left rotation. The figures show the synergy cluster centroids of these task that could be explained by either a single or linearly combined multiple synergy cluster centroids of all tasks (synergies in blue) matched by maximizing scalar product > 0.75. Observed muscle synergies extracted from the single-task EMG (orange) and their reconstructions by merging their respective W1- combinations (dark orange) were further presented.

## Discussion

Several studies have investigated shared or merged muscle synergies across different tasks such as walking and running ^8,42^, walking and cycling ^25^, various directions of reaching ^14,32^ and stepping and non-stepping postural controls ^35^. Their results indicated that different human behaviors use the fundamental motor modules that reflect the functional control units as a neural constraint on motor outputs. However, the extent to which representations of muscle synergies in the control of diverse human behaviors have not been comprehensively investigated in previous studies. In our study, we extracted muscle synergies from a large set of EMG (26 muscles) activities across bilateral locations of the trunk and lower limbs during 24 locomotion and stability tasks that were fundamental for a variety of physical activities. We found that 13 clusters of fundamental muscle synergies accounted for almost all synergy clusters of each of the 24 tasks. When we compared the synergy clusters extracted from individual tasks across participants, we found a high similarity (SP > 0.75) of a single or multiple linear combinations from the 13 fundamental muscle synergy clusters extracted from all tasks across participants. In the following sections, we discuss the possible neural mechanism underlying a diverse set of human behaviors based on the assumptions that muscle synergies represent motor modules to coordinate patterns utilized by the CNS ^24^.

### Characteristics of muscle synergies across 24 tasks

We applied cluster analysis to the muscle synergies from the all-task EMG matrix across participants, and identified 13 synergy clusters. As shown in Table 2, we broadly categorized muscle synergies into three sets based on the major contributions of the muscles (i.e., right muscle patterns, left muscle patterns, and bilateral muscle patterns). In the right and left muscle patterns, W1 and W6 were dominated by muscles around the ankle and knee joints (i.e., TA, RF, and VM). W2 and W7 were mainly composed of muscles related to the knee and hip joints (i.e., RF, VM, Gmed, and GM), and W3 and W8 employed the ankle and hip joints (i.e., MG and Gmed). Furthermore, BF mainly contributed to W4 and W9. While all four pairs were predominantly composed of extensor muscles that can move and stabilize the body during locomotion and postural tasks, they may have a distinct functional feature because the different tasks require different combinations of muscle synergies (Supplementary Table S2 online). In contrast, the pairs of W5 and W10, W11, W12, and W13 were composed of back muscles (i.e., ES, LD) and abdominal muscles (i.e., RAS and OE) either in unilateral or bilateral patterns (Table 2). Notably, they were widely observed across 24 tasks (Supplementary Table S2 online) and may be used for bilateral trunk movements or stabilization of the body accompanied by W1 to W10 with relatively low levels of trunk muscle activities when the lower limbs are moving ^43^. Although we still do not know how muscle synergies in our study arise and whether they reflect neural structure for motor outputs, 13 muscle synergies extracted from our study may form a repertoire of whole lower limb and trunk muscle activation patterns, which can be shaped by biomechanical interactions and constrain the environment through a lifetime ^18,44^.

### Hypothetical neural mechanisms underlying muscle-synergy controlling diverse behavior

If we assume that the muscle synergy extracted from the whole-task EMG matrices in our data may have a unique set of networks in which each synergy provides functionally necessary compositions in muscle activities, then one can expect that any combinations of these synergies may provide stable and predictable motor outputs in a diverse range of human behaviors ^44^. The strength of our finding is that it indicates that there is a set of fundamental muscle synergies shared with different combinations of these synergies in single and/or merging states to produce 24 locomotion and stability tasks. Considering that several previous studies in animal and human experiments have confirmed that muscle synergies observed in motor behaviors have cortical and subcortical neural underpinnings ^45–48^, it can be reasonably assumed that they are inherently robust and may be encoded in the CNS. Here, we hypothesize the existence of neural mechanisms underlying the flexible recruitment of muscle synergies in various combinations for executing a variety of movements and postures. For example, we extracted four synergy clusters in locomotion tasks, including walking, running, and bilateral jumps. Surprisingly, as shown in Fig. 2 and Supplementary Table S2 online, all synergies in the three tasks used almost the same synergies of all tasks with different combinations to be merged (SP > 0.8). Furthermore, even in the unique dynamic tasks such as cat-and-dog as well as simple axial tasks such as forward bend and rotation, subsets of these 13 fundamental muscle synergies were used, either independently or in a merging state (Fig. 3 and Supplementary Table S2 online).

Interestingly, we found that muscle synergies in 24 locomotion and stability tasks were predominantly reconstructed by merging various combinations of fundamental muscle synergies (Table 3). A study reported that muscle synergies of cycling can result from merging synergies of walking ^25^. Another recent study showed the merging of original muscle synergies during running through running training ^18^. It is suggested that merged synergies were the result of the co-recruitment of multiple muscle synergies by neural networks driving the muscle synergies represented as *C* in equation 1 ^24,44^. Based on previous studies, we speculate that the upstream driving layer (e.g., C_task_ in Fig.4) may flexibly recruit the fundamental muscle synergies (e.g., W’ in Fig. 4) located at different levels from the driving layers in the motor hierarchy to execute highly variable tasks (the schematic structure in Fig. 4). Our hypothesis is possibly equivalent to a generalized two-level CPG model for the control of locomotor muscle activity ^49^. The model consists of two distinct neural network layers: 1) a pattern formation (PF) network layer that defines groups of synergistic and antagonistic motoneuron pools and 2) a rhythm generation layer that controls the activity of PF networks. However, it should be noted that the exact neural substrates encoding muscle synergies and their driving networks in humans remain largely unknown.

**Figure 4.**
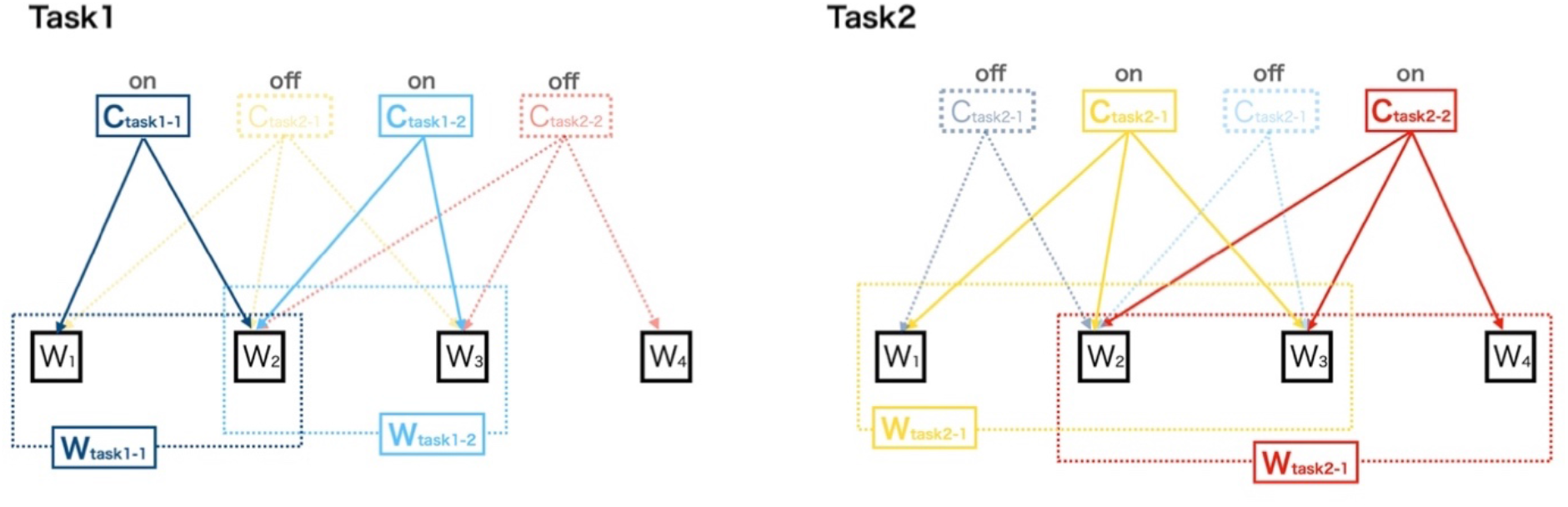
A hypothetical neural mechanism of merged fundamental muscle synergies with its temporal patterns in a diverse range of human behaviors. This model shows that the CNS flexibly recruits multiple synergies for different tasks. For example, C_task1-1_ with W_1_ and W_2_ and C_task1-2_ with W_2_ and W_3_ turn on while C_task2-1_ and C_task2-2_ turn off to execute task 1. Similarly, C_task1-1_ with W_1_, W_2_, and W_3_ and C_task1-2_ with W_2_, W_3_, and W_4_ turn on, ceasing to be active in C_task1-1_ and C_task1-2_ for task 2.

Since we propose that upstream driver *C* presents synchronous recruitments of the fundamental muscle synergies that have distinct functional roles in organizing muscle synergies for the 24 locomotion and stability tasks, it is possible that the CNS may also coordinate other simple or complex human behaviors using certain combinations of these synergies. Thus, muscle synergies during human behaviors found in previous extensive research may reflect layered structures composed of the fundamental muscle synergies extracted from our study. The advantage of these hypothetical mechanisms is that it prevents the sum of all muscle synergies from exceeding the number of relevant muscles utilized during diverse human behaviors, supporting the premise of compendium in coordinative patterns to execute several movements under different biomechanical conditions ^23^. Further research is needed to investigate the muscle synergies identified by factorization algorithms coupled with CNS manipulations and/or neural recordings (e.g., CNS stimulations, spinalization, and electroencephalogram) to validate the neural representation of the fundamental muscle synergies observed in our study ^24^.

### Clinical implications

The results of this study may have several clinical implications. First, several studies have investigated muscle synergies in individuals with different characteristics, such as musculoskeletal and neurological disorders ^50–52^ as well as athletes ^8,18,53^. Since we identified the fundamental muscle synergies that may underlie diverse human behaviors in healthy individuals, investigating the changes in muscle synergies such as the number of synergies as well as their compositions in a population of interest may facilitate the understanding of distinct features in motor controls that are associated with severity of symptoms ^50,54^ or that profile myriad skills and performance in athletes ^27,28^. Second, recent studies have shown the efficacy of muscle synergy-based interventions using functional electrical stimulations (FES) on motor performance in stroke survivors ^55,56^. Since our study found that each synergy may have functionally plausible patterns that play an important role in executing diverse human movements and postures, it may provide a rationale for designing interventions that use FES to focus on these functional sets of muscle synergies to improve motor performance. Lastly, we found that different tasks with various biomechanical demands and constraints may largely share the same muscle synergies with different combinations of synergies to be merged. Thus, clinicians may choose to intensively train a particular task to transfer the effectiveness to other tasks ^57^, given that the transfer of motor learning effects among tasks will be high when muscle synergies involved in different motor tasks are shared ^58^.

### Limitations

Our study had several limitations. First, it has been reported that the number of recording muscles may affect the amount and structure of muscle synergies ^59^. Although EMG recordings in our study were relatively large (i.e., 26 EMG channels), we limited the recording of EMGs from only the major muscles in the trunk and lower limbs. Similarly, we were also limited to 24 fundamental tasks that involved only locomotion and postural tasks. As such, tasks that accompany coordination between the upper limbs, trunk, and lower limbs were not considered ^5^. Thus, it is conceivable that some relevant muscle synergies may have been missed in our study. Second, because we used a larger set of EMG recordings and tasks, our time constraint during experiments precluded the measurement of kinematic data such as joint angles as well as velocities, and allowed variability of movements in each task, which may impact muscle synergy extractions. The lack of availability of kinematic data ceases to separates the movement phase and thus unable to investigate the contributions of the fundamental muscle synergies for each phase of each task ^44^. Lastly, although we extract the fundamental muscle synergies using NMF that may present neural mechanisms for diverse human behaviors, whether the factorization-derived synergies reflect neural organization to coordinate human behaviors remains questionable ^24^. This can be due to the possibility that extracted muscle synergies represent biomechanical constraints of tasks rather than neural constraints ^60^ and the nonlinearity in magnitude summations of the EMG or force vectors ^61,62^.

## Conclusion

In this paper, we extracted a repertoire of fundamental muscle synergies from the EMGs during a variety of human behaviors that involve trunk and lower limb movements in healthy individuals. We found that the flexible recruitment of the fundamental muscle synergies in either the independent or merging state can account for almost all 24 behaviors, including locomotion and stability tasks. Our findings may support the notion that low dimensional motor modules are required in a diverse range of human behaviors with different biomechanical contexts.

## Supporting information

10.5281/zenodo.5119976

## Data availability

All data are available upon reasonable request.

## Acknowledgements

We would like to thank Editage (www.editage.com) for English language editing.

## Author contributions

Conception and design of the work H.S., H.Y. and A.S.; data acquisition H.S., A.S., and T.K.; data analysis H.S. and H.Y.; data interpretation H.S., H.Y., A.S., T.K. and K.N.; drafted the manuscript H.S., H.Y., A.S., T.K. and K.N. All authors approved the final version of the manuscript and agree to be accountable for the content of the work.

## Additional Information

The authors declare no competing interests.

