## Supplementary material for "Flexible recruitments of fundamental muscle synergies in the trunk and lower limbs for highly variable movements and postures": 10.5281/zenodo.5119976

**This file includes:**

Supplementary Figures S1 and S2

Supplementary Tables S1 and S2

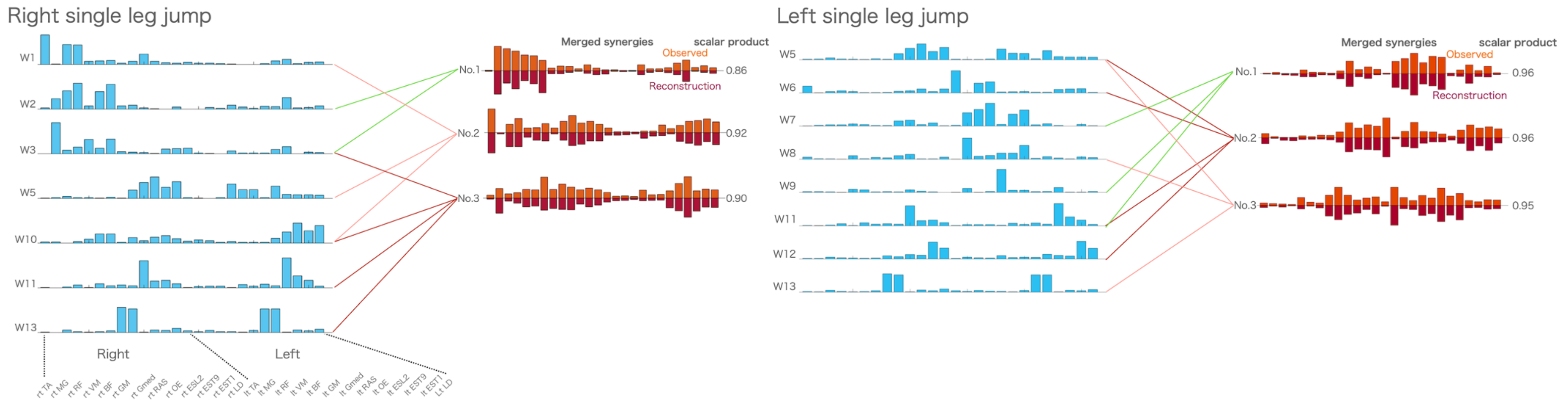

**Supplementary Figure S1. Relationship between muscle synergies of all tasks and muscle synergies of other locomotion tasks.** Shown are synergy cluster centroids of these task that could be explained by either a single or linearly combining multiple synergy cluster centroids of all tasks (synergies in blue) matched by maximizing scalar product > 0.75. Observed muscle synergies extracted from the single-task EMG (orange) and their reconstructions by merging their respective W1- combinations (dark orange) were presented.

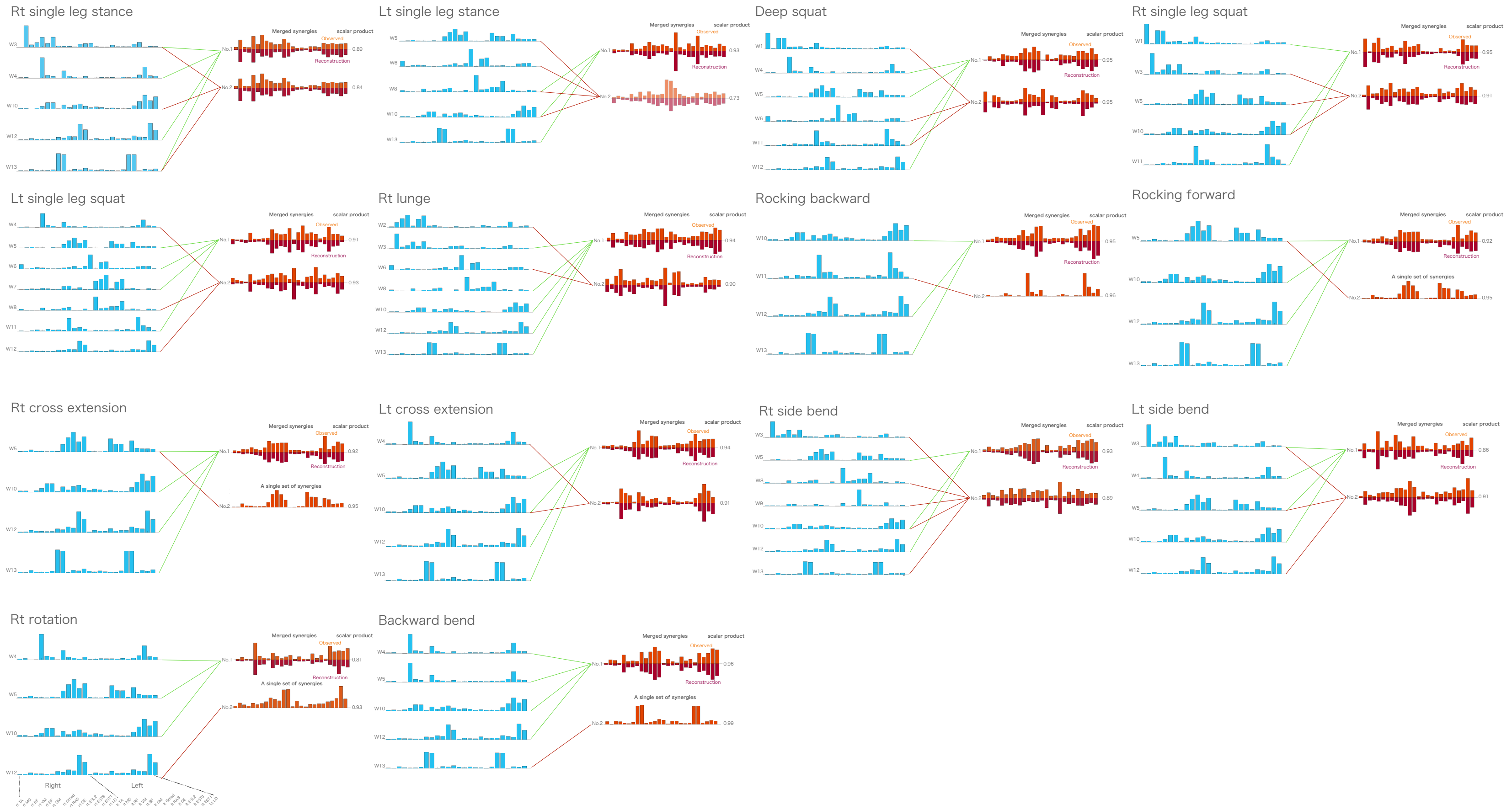

**Supplementary Figure S2. Relationship between muscle synergies of all tasks and muscle synergies of other stability tasks.** Shown are synergy cluster centroids of these task that could be explained by either a single or linearly combining multiple synergy cluster centroids of all tasks (synergies in blue) matched by maximizing scalar product > 0.75. Observed muscle synergies extracted from the single-task EMG (orange) and their reconstructions by merging their respective W1- combinations (dark orange) were presented.

**Supplementary Table S1: Full descriptions of movement and postural tasks.**

| Categories | | Task numbers | Tasks | Repetitions, duration | Conditions |
|---|---|---|---|---|---|
| Locomotion | | 1 | Walk (1.5 m/s) | 30 seconds | Walked on a treadmill (Bertec, Columbus, OH, USA) at 1.5m/s |
| | | 2 | Run (2.7 m/s) | 30 seconds | Run on a treadmill (Bertec, Columbus, OH, USA) at 2.7m/s |
| | | 3 | Bilateral jump | 8 repetitions | From a standing position, jump upward with arms freely in both sides and to maintain the same position at the instants of take-off and landing. After that, return to a standing position |
| | | 4 | Rt single leg jump | 8 repetitions | From standing on a right leg, jump upward with arms freely in both sides and to maintain the same position at the instants of take-off and landing |
| | | 5 | Lt single leg jump | 8 repetitions | From standing on a left leg, jump upward with arms freely in both sides and to maintain the same position at the instants of take-off and landing |
| | | 6 | Sit to stand to sit | 8 repetitions | Sit to stand to sit from a chair (40 cm height) |
| Stability | Static postures | 7 | Rt single leg stance | 15 seconds | Single leg standing on a right leg |
| | | 8 | Lt single leg stance | 15 seconds | Single leg standing on a left leg |
| | Dynamic postures | 9 | Deep squat | 8 repetitions | From a standing position with hands raised, squatting approximately 120 degree of knee flexion and return |
| | | 10 | Rt single leg squat | 8 repetitions | From a standing position on a right leg with arms freely in both sides, squatting approximately 100 degree of knee flexion and return |

| | | | | | |
|---|---|---|---|---|---|
| | | 11 | Lt single leg squat | 8 repetitions | From a standing position on a left leg with arms freely in both sides, squatting approximately 100 degree of knee flexion and return |
| | | 12 | Rt lunge | 8 repetitions | From a standing position with a right leg forward, lowering a body until a left knee almost touches a floor and return |
| | | 13 | Lt lunge | 8 repetitions | From a standing position with a left leg forward, lowering a body until a right knee almost touches a floor and return |
| | | 14 | Rocking backward | 8 repetitions | In a quadruped position, transfer of the buttock backwards ("rocking") keeping low back in neutral until knees reach approximately 130 degree of flexion and return |
| | | 15 | Rocking forward | 8 repetitions | In a quadruped position, transfer of the buttock forward ("rocking") keeping low back in neutral until knees reach 0 degree of hip extension and return |
| | | 16 | Rt cross extension | 8 repetitions | In a quadruped position, raise a right arm and a left leg straight out and return |
| | | 17 | Lt cross extension | 8 repetitions | In a quadruped position, raise a left arm and a right leg straight out and return |
| | | 18 | Cat-and-dog | 8 repetitions | In a quadruped position, round a back and drop a chin to a chest (cat) and then lift a head up and arch a back down toward a floor (dog). After that, return to a quadruped position |
| | Axial | 19 | Forward bend | 8 repetitions | From a standing position, bend a trunk forward as far as possible and return |
| | | 20 | Rt side bend | 8 repetitions | From a standing position, bend a trunk to a right side as far as possible and return |

| | | | | | |
|---|---|---|---|---|---|
| | | 21 | Lt side bend | 8 repetitions | From a standing position, bend a trunk to a left side as far as possible and return |
| | | 22 | Backward bend | 8 repetitions | From a standing position with arms raised, bend a trunk backward as far as possible and return |
| | | 23 | Rt rotation | 8 repetitions | From a standing position, rotate a trunk to a right side as far as possible and return |
| | | 24 | Lt rotation | 8 repetitions | From a standing position, rotate a trunk to a left side as far as possible and return |

Each participants performed 24 locomotion and stability tasks. The order of tasks was randomly assigned.

**Supplementary Table S2: Contributions of 13 synergy clusters of all tasks for each task execution.**

| | | | | | | | | Static postures | | Dynamic postures | | | | | | | | | | Axial | | | | | |
|---|---|---|---|---|---|---|---|---|---|---|---|---|---|---|---|---|---|---|---|---|---|---|---|---|---|
| | | Walk | Run | BilJP | Rt SJP | Lt SJP | STS | Rt SLS | Lt SLS | DS | Rt SS | Lt SS | Rt LG | Lt LG | RB | RF | Rt CE | Lt CE | CD | FB | Rt SB | Lt SB | BB | Rt RT | Lt RT |
| Right patterns | W1 | ✓ | | ✓ | ✓ | | ✓ | | | ✓ | ✓ | | | ✓ | | | | | | | | | | | |
| | W2 | ✓ | ✓ | ✓ | ✓ | | | | | | | | ✓ | | | | | | | | | | | | |
| | W3 | ✓ | ✓ | ✓ | ✓ | | | ✓ | | | ✓ | | ✓ | ✓ | | | | | | ✓ | ✓ | ✓ | | | |
| | W4 | ✓ | ✓ | | | | | ✓ | | ✓ | | ✓ | | | | | | ✓ | | ✓ | | ✓ | ✓ | | ✓ |
| | W5 | ✓ | ✓ | ✓ | ✓ | ✓ | ✓ | | ✓ | ✓ | ✓ | ✓ | | ✓ | | ✓ | ✓ | ✓ | | ✓ | ✓ | ✓ | ✓ | ✓ | ✓ |
| Left patterns | W6 | ✓ | ✓ | ✓ | | ✓ | ✓ | | ✓ | ✓ | | ✓ | ✓ | ✓ | | | | | | | | | | | |
| | W7 | ✓ | ✓ | ✓ | | ✓ | | | | | | ✓ | | ✓ | | | | | | | | | | | |
| | W8 | ✓ | ✓ | ✓ | | ✓ | | | ✓ | | | ✓ | ✓ | ✓ | | | | | | | ✓ | | | | |
| | W9 | | ✓ | | | ✓ | | | | | | | | | | | ✓ | | | ✓ | ✓ | | | | |
| | W10 | ✓ | ✓ | ✓ | ✓ | | | ✓ | ✓ | | ✓ | | ✓ | ✓ | ✓ | ✓ | ✓ | ✓ | ✓ | ✓ | ✓ | ✓ | ✓ | ✓ | ✓ |
| Bilateral patterns | W11 | ✓ | ✓ | ✓ | ✓ | ✓ | ✓ | | | ✓ | ✓ | ✓ | | | | | ✓ | | ✓ | ✓ | | | | | |
| | W12 | | ✓ | ✓ | | ✓ | ✓ | ✓ | | ✓ | | ✓ | ✓ | ✓ | ✓ | ✓ | ✓ | ✓ | ✓ | ✓ | ✓ | ✓ | ✓ | ✓ | ✓ |

| | | | | | | | | | | | | | | | | | | | | | |
|---|---|---|---|---|---|---|---|---|---|---|---|---|---|---|---|---|---|---|---|---|---|
| | W13 | ✓ | ✓ | ✓ | ✓ | ✓ | | ✓ | ✓ | | | | ✓ | | ✓ | ✓ | ✓ | ✓ | ✓ | ✓ | ✓ |

Shown are the contributions of each 13-synergy cluster of all tasks in single or linear combination of these synergies for each task execution (Scalar product > 0.75). Bil: bilateral; Rt: right; Lt left; JP: jump; SJP: single leg jump; STS: sit-to-stand-to-sit; SLS: single leg stance; DS: deep squat; SS: single leg squat; LG: lunge; RB: rocking backward; RF: rocking forward; CE: cross extension; CD: cat-and-dog; FB; forward bend; SB: side bend; BB: backward bend; RT: rotation.
